# Sustained 10% Oxygen Promotes Atrial Rather Than Ventricular Specification During Human iPSC-Cardiomyocyte Differentiation

**DOI:** 10.64898/2026.09.16.750048

**Authors:** Sabrina Bech Mathiesen, Frederik Adam Bjerre, Anne Kathrine Søgaard Terp, Ditte Gry Ellman, Jannik Hjortshøj Larsen, Peer Bendix Horn, Per Svenningsen, Ellen Ngar Yun Poon, Charlotte Harken Jensen, Ditte Caroline Andersen

## Abstract

**Background:** Induced pluripotent stem cell-derived ventricular cardiomyocytes (iPSC-vCMs) hold great promise for replacing ventricular cardiomyocytes lost after myocardial infarction. However, their immature phenotype limits successful engraftment by increasing the risk of post-transplant arrhythmias. In contrast to the atmospheric O_2_ used during most iPSC-vCM differentiations, O_2_ levels inside the developing heart remain low, but very little is known on the effect on O_2_ on iPSC-vCM differentiation.

**Methods:** We used a GMP compliant Quad Physoxia glovebox platform providing continuous stable specified O_2_ tensions during all processes to simulate the *in vivo* O_2_ conditions more closely with the aim of improving iPSC-vCM maturation.

**Results:** We demonstrate by single cell RNA sequencing, and data integration with datasets for human cardiomyocytes from the different heart chambers as well as cell morphology-, ploidy-, and functional studies, that sustained 10% O_2_ throughout iPSC-CM differentiation promotes atrial-instead of ventricular iPSC-CM subtype specification.

**Conclusions:** While this rejects our original hypothesis and forces some concerns to iPSC-vCM manufacturing by sphere technology, these unexpected data may serve as an attractive and easy approach to refine atrial iPSC-CM specification to benefit their exponentially growing diagnostic-, cytotoxic-, and regenerative use.

## Introduction

It is generally known that the human heart is unable to repair damaged ventricular cardiomyocytes (vCM) after myocardial infarction (MI), which may lead to heart failure and ultimately death(1). Except for heart transplantations, no curative treatment exists, but induced pluripotent stem cell-derived vCM (iPSC-vCM) remains a virtually inexhaustible source for cardiomyocyte (CM) replacement therapy(2). However, among other challenges iPSC-vCM immaturity(3) causes arrythmia upon intracardiac transplantation and limits their translation(4–7). In most studies, iPSC-vCM are differentiated at atmospheric O_2_(_4-9_), and at their best corresponds to the phenotype of a fetal vCM(3). However, considering heart development, vCM before birth reside in a relatively hypoxic (25-30 mmHg; ∼3-5% O_2_) environment(10, 11), which then rises after birth to 75-100 mmHg; ∼11-15% O_2_(11). Still, this is far below the atmospheric (160 mmHg; ∼21%) O_2_ tension generally used during *in vitro* iPSC-vCM-derivation. The vast increase in blood oxygenation around birth is known to correlate with a switch in energy metabolism from glycolysis to oxidative phosphorylation and ultimately CM cell cycle arrest(12) with subsequent polyploidization(3, 13, 14), all important events in terminal vCM differentiation and maturation(3, 15). Yet very little is known about the effect of O_2_ on vCM differentiation from pluripotent stem cells (16–22). A few studies have suggested that lower O_2_ tensions increase the vCM yield(18–21), whereas two other studies report an inhibitory effect on vCM differentiation(16, 20). These studies exploit "low" O_2_ (typically 2-5%) for only a short period (24-48 hours)(20, 23) or only during the first 9-16 days(18, 19, 21–23) of differentiation and several of the studies rely solely on embryoid body formation based vCM differentiation protocols(18, 19, 21), whereas none of the studies handle the cells at continuous O_2_ tensions at all times, which may be critical for stem cells(24).

Herein, we therefore specifically hypothesized: that the most used small molecule-based iPSC-vCM specification approach, combined with tightly controlled physiological low O_2_ tensions throughout vCM differentiation (until day 30), including a shift at day 12 to 10% O_2_ tension simulating birth, would enhance iPSC-CM journey towards their mature adult vCM phenotype as compared to those obtained at atmospheric O_2_ tension conditions(25)

## Results

### Single cell transcriptomic profiling reflects an iPSC-CM phenotype change with "low" physiological O_2_ tensions

To test our hypothesis, we employed an iPSC-line with a mono-allelic mEGFP tag at the c-term of the myosin light chain 2 ventricle isoform (MYL2 or MLC2V), a marker of vCM. We refined a previously tailored protocol(26) based on modulation of regulators of the Wnt pathway(17, 27) and metabolic selection(28) to the iPSC-line (Figure 1a). At first, we confirmed robustness of our CM differentiation protocol at atmospheric O_2_. Already, at differentiation day 4-8 (D4-8), gene expression analysis showed a marked decrease in pluripotency (Figure 1b) with a concomitant emergence of CM markers (Figure 1b). After metabolic selection (Figure 1a), ventricle- and maturation markers increased substantially (Figure 1b+c), and at D30, flow cytometry (Figure 1d) verified iPSC-CM homogeneity with 98.5%±1.3% being TROPONIN T^+^ CM (Figure 1e) and 67.4%±6.5% of these were of the ventricular CM subtype (Figure 1f). Cardiac lineage commitment was further supported by widespread protein immunofluorescence analysis of the cardiac/progenitor transcription factors GATA-4 and NKX-2.5 (Figure 1g, top panel), the pan cardiac markers of TROPONIN T, TROPOMYOSIN and ACTININ (Figure 1g, middle and lower panel), and the vCM marker MLC2v (Figure 1g, lower panel). Using this iPSC-vCM protocol, we next mimicked the *in vivo* scenario of heart development with physiological "low" O_2_ tension (2.5-10%) during cardiac specification followed by 10% O_2_ the remaining period simulating cardiac development after birth (Figure 1h). Previous studies have suggested that even short durations of atmospheric O_2_ alter stem cell gene expression(24). Thus, to tightly control the O_2_ tension during iPSC-vCM derivation, we exploited a GMP compliant SCI-TIVE Quad Physoxia glovebox platform providing continuous stable, specified O_2_ tensions during all processes (Figures 1i and S1). The cells are thus never exposed to ambient air unless in the 21% O_2_ group. Initially, we equilibrated and tested iPSCs at 10% O_2_. Gene expression analysis of markers for pluripotency (*OCT4, NANOG, SOX2*) and hypoxia (*HIF-a*) suggested that the abrupt change in O_2_ (21-versus 10%) did not induce any immediate consistent and damaging changes in iPSCs during expansion (Figure S2a-b). We next used 2.5-10% O_2_ for 12 days followed by 10% O_2_ (D10-D30) comparing to those derived at continuous 21% O_2_ for the entire period and performed serial derivations of iPSC-CM with subsequent analysis at D30. Flow cytometry (Figure 1j) for Troponin T^+^ showed that cardiac specification was consistently high (2.5% O_2_: 98.8%±1.3%; 5% O_2_: 98.2±0.2%; 10% O_2_: 96.6%±1.3%; 21% O_2_: 99.4%±0.5%) across all O_2_ levels (Figure 1k) despite that overall appearance and viability (data not shown) seemed slightly compromised at 2.5- and to some extent at 5% O_2_ tensions. The latter being in agreement with a previous report(22). Then to investigate how the difference in O_2_ tension affected iPSC-vCM maturation, we performed single cell RNA sequencing (scRNA-seq). All quality parameters of the scRNA-seq running(29) were high (Figure S3a-d) validating the scRNA-seq data quality. In total, we obtained 28,689 cells, which were then filtered for CM markers (*ACTC1, TNNT2, TNNC1*) (Figure S3e). These data confirmed an overall high (96-99%) iPSC-CM purity, albeit only around 80% passed the triple filter for the iPSC-CM derived at 5% O_2_ (Figures 1l and S3e-h). Based on gene expression: profiling, re-integration, clustering, and visualization of the filtered iPSC-CM were then performed using Uniform Manifold Approximation and Projection (UMAP) plotting (Figure 1m+n). iPSC-CM cultured at 2.5-10% O_2_ clustered together, yet remarkably distinct from iPSC-CM at 21% O_2_ suggesting a phenotype switch upon O_2_-lowering, despite their similar CM specification (Figures 1k+l and S3e-h).

**Figure 1.**
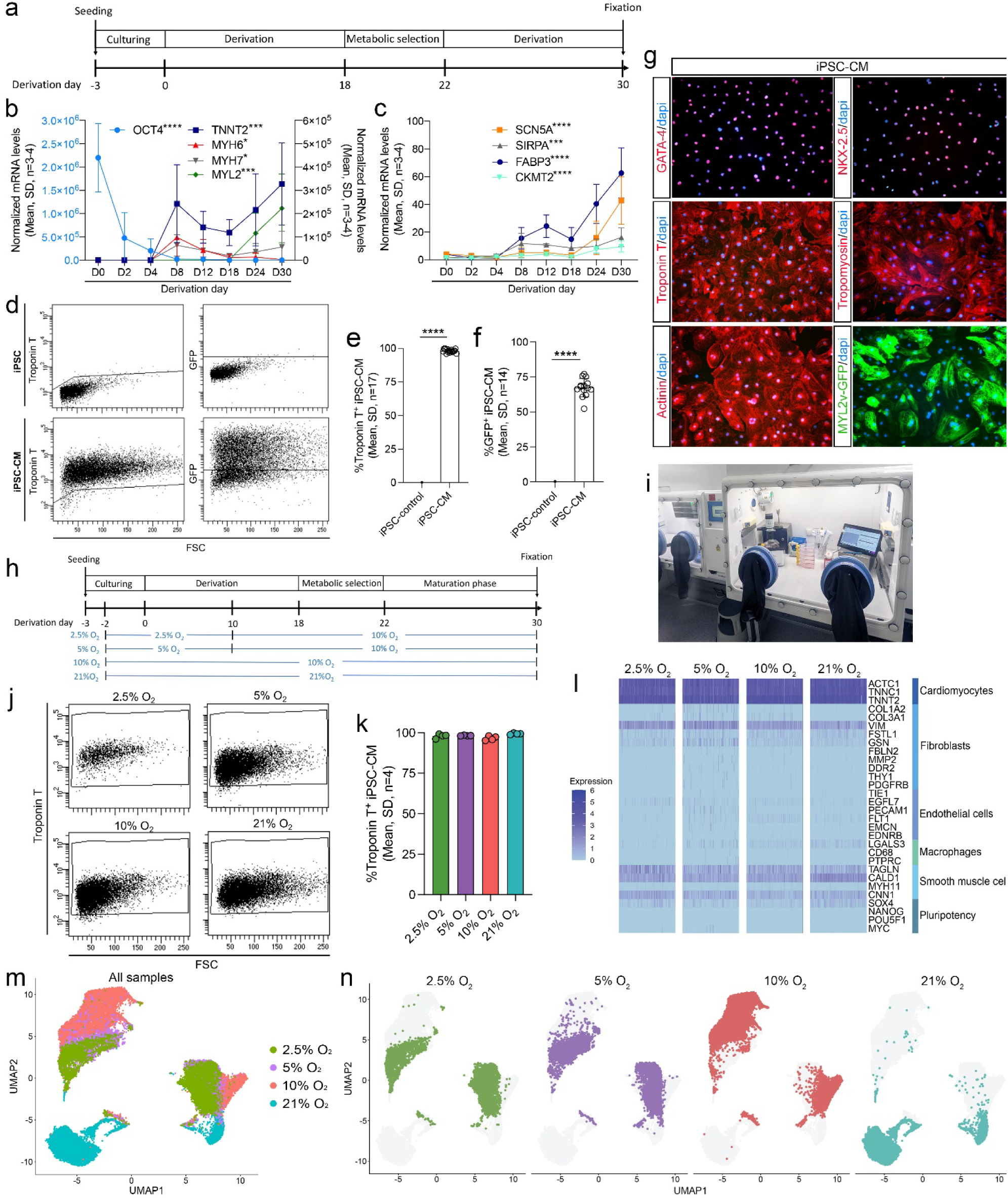
iPSC-CM single cell transcriptomic patterns change with O2 tensions during derivations. **a**, Schematic view of the iPSC-CM derivation schedule from iPSCs expressing a mEGFP tag at the c-term of the ventricular marker myosin light chain 2 ventricle isoform (MYL2/MLC2v). **b-c**, Normalized (GAPDH and B2M) mRNA levels analyzed by qRT-PCR of **b**, OCT4 (pluripotency marker), TNNT2, MYH6, MYH7, MYL2 (CM and ventricular markers) **c**, SCN5A, FABP3, SIRPA and CKMT2 (CM maturation markers) at indicated derivation day (D). Mean, SD, n = 4. Statistical analysis included ordinary, one-way ANOVA. \**P*≤0.05, \*\*\**P*≤0.001, \*\*\*\**P*≤0.0001. **d**, Representative flow cytometric dot plots for Troponin T-(left) and GFP^+^ (right) expression in iPSC-CM at D30 (Bottom) as compared to an iPSC control group (Top). **e-f**, Flow cytometric quantification of %Troponin T and %GFP^+^ in iPSC-CM at D30. Mean, SD, n = 17. **g**, Immunocytochemistry of D30 iPSC-CM (DAPI (nuclei) in blue; GATA-4, NKX-2.5, Troponin T, Tropomyosin, Actinin in red; MYL2v-GFP in green). **h**, Schematic overview of the iPSC-CM derivation protocol at different O_2_ tensions (2.5 to 10%, 5 to 10%, continuous 10% or 21% O_2_) as controlled using **i**, pre-equilibrated SCI-TIVE Quad Physoxia closed glovebox workbenches (GMP facility/OUH-CELL-BENCH). **j**, Representative flow cytometric dot plots of Troponin T^+^ D30 iPSC-CM derived at the four different O_2_ tensions (2.5 to 10%, 5 to 10%, continuous 10% or 21% O_2_). **k**, Flow cytometric quantification of %Troponin T^+^ D30 iPSC-CM as depicted in j. Mean, SD, n = 4. **l**, Heatmap visualizing normalized expression of known marker genes for different cell types (CM, fibroblasts, endothelial cells, macrophages, smooth muscle cells and pluripotent cells) in scRNA-seq data of D30 iPSC-CM derived at the four O_2_ conditions (2.5 to 10%, 5 to 10%, continuous 10% or 21% O_2_). **m**, UMAP embeddings showing the D30 iPSC-CM scRNA-seq data as merged or **n**, as individually depiction of iPSC-CM derived at 2.5 to 10%, 5 to 10%, continuous 10% or 21% O_2,_ respectively.

### Physiological "low" O_2_ tension drives an atrial-like cardiomyocyte iPSC lineage transcriptomic signature

Then to avoid any interference from potential low viability cells in 2.5-5% O_2_ groups and to investigate the observed phenotype difference, we generated a new dataset for 10- and 21% O_2_ iPSC-CM (Figure 2a) and performed Louvain clustering resulting in six different clusters (1–6) (Figure 2b). Combining UMAP and Louvain clustering analysis, we found that cluster 2+3 and 1 were specific for 10- and 21% O_2_ iPSC-CM, respectively, whereas cluster 4-6 were shared between them (Figure 2a+b). All clusters expressed CM genes (*TNNT2, TNNC1, MYL7, ACTC1*) as expected (Figure 2c). Cluster 1 and 5 represent ventricular (positive for: *MYL2, MYH7*) iPSC-CM, while cluster 2 and -3 showed a more atrial like (positive for *TBX5, CACNA1D, KCNJ3* and negative for *MYL2, MYH7*) CM subtype (Figure 2c). Indeed, ∼50% of the iPSC-CM at 21% O_2_ embraced a clear ventricular signature, while ∼70% of iPSC-CM at 10% O_2_ could be classified as atrial-like cells (Figure 2d). The very small cluster 6 was more diffuse expressing both atrial and ventricular genes, while cluster 4 had limited expression of specification and thus seemed to represent more immature CM (Figure 2c). These data thus surprisingly suggested that physiological lower levels like 10% O_2_ during iPSC-CM derivation somehow favors atrial specification at the expense of the vCM phenotype. Initially, we speculated whether this was a simple matter of halted differentiation of the iPSC-vCM at 10% O_2_. Yet, scRNA-seq of immature D11 iPSC-vCM derived at 21% O_2_ did not share this atrial-like signature (Figure 2e) demonstrating that atrial differentiation seemed specific to the 10% O_2_ derivation schedule. Moreover, we found that our atrial-like iPSC-CM derived at 10% O_2_ were *GJC1*-(*CX45*), *HCN4*-, *TBX3*-, *TBX18*-, *GJA1*+ (*CX43*), *NKX-2.5*+ (Figure S4), which excluded that they were of the pacemaker(30, 31) CM subtype.

**Figure 2.**
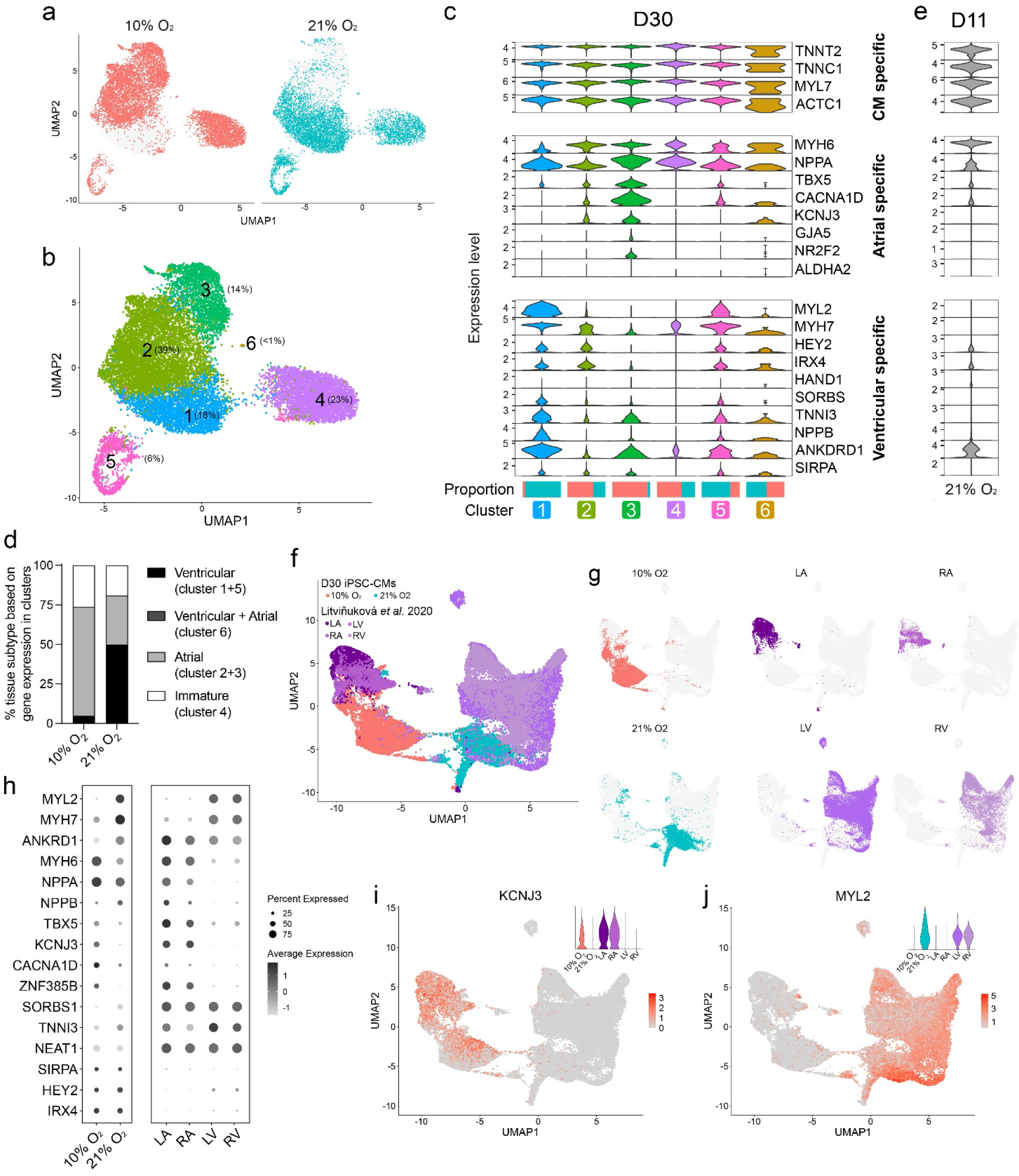
Physiological "low" O2 tension drives an atrial cardiomyocyte iPSC transcriptomic signature. **a**, UMAP-plots of iPSC-CM split by cells derived at 10-(left) and 21% O_2_ (right). **b**, UMAP-plots colored by Louvain clustering based on both 10- and 21% O_2-_derived iPSC-CM. **c**, Violin plots visualizing normalized expression of known marker genes for general CM specification, arterial specification, ventricular maturation specification and cell cycling identifying the expressional specificity of each cluster. Proportions of 10- and 21% O_2_-derived iPSC-CM for each cluster are visualized as bar plots below. The normalized expression of selected genes is visualized in UMAP embeddings. **d**, Percentages of 10- and 21% O_2-_derived iPSC-CM belonging to each tissue subtype (ventricular, atrial and immature) based on UMAP clustering of known marker genes. **e**, Vulcano plot visualizing differential expression of known marker genes for CM specification, arterial specification, ventricular maturation specification and cell cycling for derivation day 11 (D11) iPSC-CM derived at 21% O_2_. **f**, UMAP embedding of adult human cardiomyocytes obtained from Litviňuková et al. dataset integrated with 10- and 21% O_2-_derived D30 iPSC-CM. **g**, UMAP visualization of 10- and 21% O_2_ D30 iPSC-CM and the Litviňuková et al. dataset divided into cells originating from the left atria (LA), the right atria (RA), the left ventricle (LA) and the right ventricle (RV). **h**, Dot plot of cardiomyocyte marker genes from 10- and 21% O_2_-derived D30 iPSC-CM and Litviňuková et al. dataset. **i-j**, UMAP embeddings visualizing the normalized expression of **i**, KCNJ3 and **j**, MYL2 in the integrated 10- and 21% O_2_-derived D30 iPSC-CM and Litviňuková dataset. The normalized expression is additionally shown in violin plots colored for by sample (as represented in g in the bottom left corner of both UMAP embeddings.

To better confirm this unexpected observation, we next integrated our scRNA-seq data of the iPSC-CM derived at 10- and 21% O_2_ with single nuclei RNA sequencing (snRNA-seq) data published by Litviňuková et al.(32) of adult human heart left and right ventricle (LV and RV) as well as left and right atrium (LA and RA). We hypothesized that iPSC-CM derived at 10- and 21% O_2_ should cluster with adult human atrium- and ventricle CM, respectively. Indeed, visual appearance showed that iPSC-CM derived at 10% O_2_ were more pronounced with adult human atrium CM, whereas iPSC-CM derived at 21% O_2_ showed greater similarities with adult human vCM (Figure 2g). This was supported by dot plot gene expression patterns of selected atrial- and ventricular markers (Figure 2h) and further visualized for the major atrial- and ventricular specific KCNJ3 (Figure 2i) and MYL2 (Figure 2j) genes, respectively. As expected, none of the iPSC-CM clusters overlapped completely with the adult human CM clusters, likely due to their general known iPSC-CM immaturity(4–7, 26) but possibly also due to the expected difference between scRNA-seq and snRNA-seq data(33). Still these integrative sc/nRNA-seq data comparisons evidenced that 10% O_2_ unintended directed iPSC-CM differentiation towards atrial-but not ventricular specification as we originally hypothesized herein.

### iPSC-CM differentiation is affected early by "low" O_2_, directing a CM phenotype switch with a change in -morphology, -ploidy and -functional calcium transients

Corroborating the data above and in agreement with an atrial phenotype, we found that iPSC-CM derived at 10% O_2_ were smaller in morphology than their iPSC-CM counterparts derived at 21% O_2_ (Figure 3a+b). They also expressed less MYL2 protein, the key indicator of ventricular specificity (Figure 3a+b), and were less binucleated (Figure 3c) despite they exhibited a higher level of ploidy (Figure 3d-e). By use of fluorescence-based measurements of intracellular calcium transients(34), we determined whether the 10- and 21% O_2_-derived iPSC-CM were functionally different, as one should expect from their different atrial- and ventricular CM subtypes. Both 10- and 21% O_2_-derived iPSC-CM showed spontaneous contractions and calcium transients (Figure 3f) with a similar calcium transient frequency (Figure 3g) and duration (Figure 3h). Yet, the amplitudes of the calcium transients (Figure 3i-k) and the time from baseline to peak calcium concentration (Figure 3l) were significantly higher for iPSC-CM derived at 21% O_2_. Consistent with this, the rate of increase in calcium concentration was similar between the 10- and 21% O_2_-derived iPSC-CM (Figure 3m). On the other hand, the return from peak to baseline calcium concentration was significantly faster for iPSC-CM derived at 21% O_2_ (Figure 3n+o). These spontaneous contraction and calcium transient differences overall support the hypothesis and above findings that O_2_ levels during iPSC-CM derivation are critical for specifying the CM subtype. Interestingly, previous findings by the Keller group(19) demonstrate that ventricular and atrial PSC-CM already at day 3-4 of differentiation are specified from distinct GYPA+/CYP261A+ and ALDH1A2+ mesodermal progenitors, respectively. Herein, we found that *MESP1* (Figure 3p) was substantially increased in 10-versus 21% O_2_-derived iPSC-CM already at D3, and this indeed coincides with increased *ALDH1A2* expression (Figure 3p) further suggesting that the CM subtype fate towards an atrial fate is initiated early during mesodermal specification and may persist and be further enhanced by our continuous low O_2_ approach.

**Figure 3.**
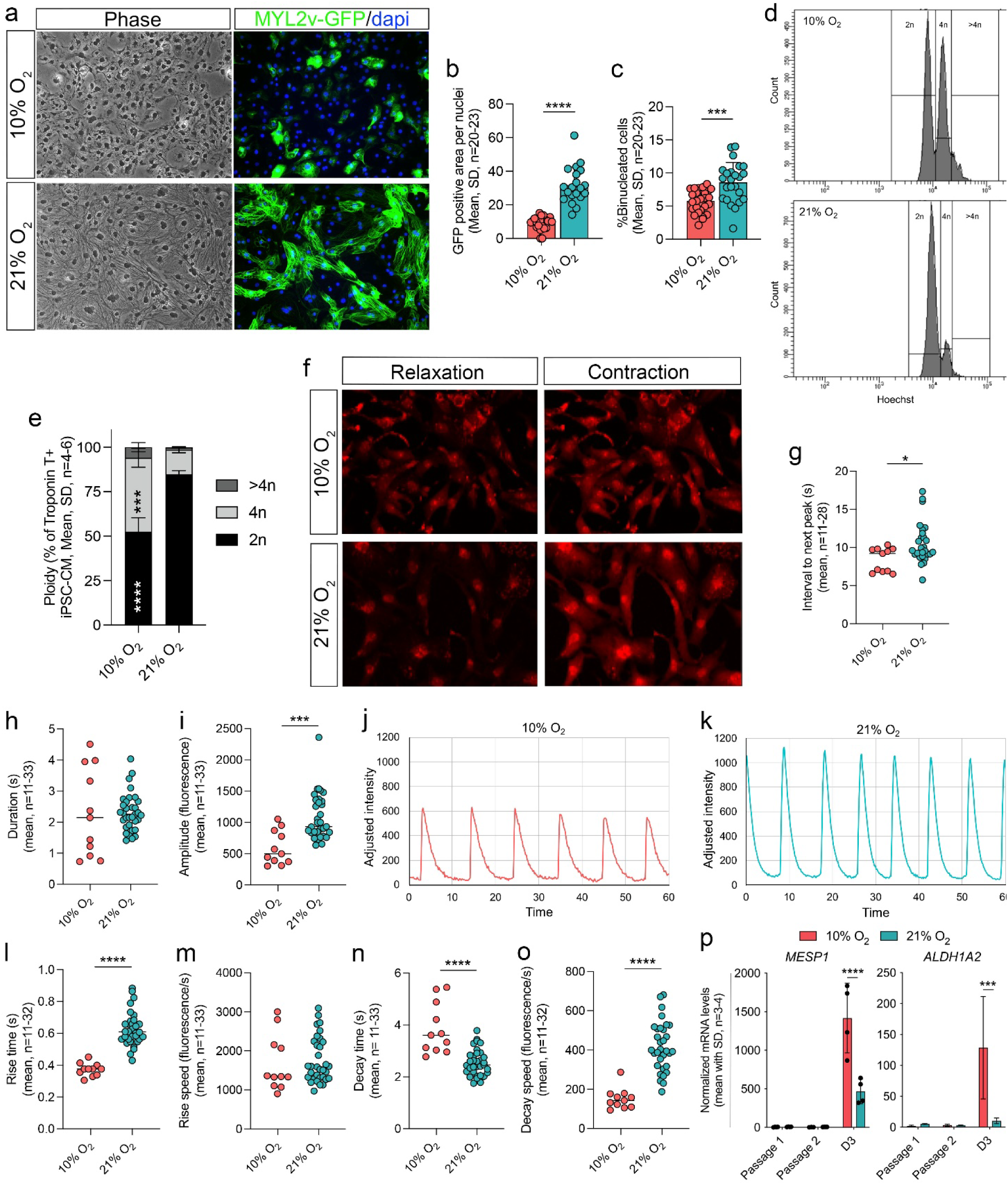
iPSC-CM derived at 10% O2 exhibit structural and functional phenotypic characteristics different from ventricular iPSC-CM. **a**, Immunocytochemistry of derivation day 30 (D30) iPSC-CM derived at 10% O_2_ (left) and 21% O_2_ (right) (phase (top); DAPI in blue/MYL2v-GFP in green, (bottom)). **b**, GFP positive area per nuclei of iPSC-CM derived at 10- and 21% O_2_ quantified from immunocytochemistry data. Mean, SD, n = 20-23. Statistical analysis included unpaired t-test; \*\*\*\**P*≤0.0001. **c**, Percentages binucleated cells derived at 10- and 21% O_2_ quantified from immunocytochemistry data. Mean, SD, n = 20-23. Statistical analysis included unpaired t-test; \*\*\**P*≤0.001. **d**, Ploidy of troponin T positive D30 iPSC-CM derived at 10- and 21% O_2_ **e**, quantified based on Hoechst intensity by flow cytometry. Mean, SD, n = 4-6. **f-o**, Optical calcium (Rhod-2 AM) confocal imaging and calcium recordings of D30 iPSC-CM derived at 10- and 21% O_2_ (n = 11 for 10% O_2_ iPSC-CM and n = 33 for 21% O_2_ iPSC-CM). **f**, Depiction of contracting iPSC-CM derived at 10-(Top) and 21% O_2_ (Bottom) at relaxation (Left) and contraction (Right) from calcium recordings. **g**, Interval to next peak (Peak to peak, s). **h,** Calcium transients duration (s). **i**, Amplitude of calcium induced fluorescence. **j**, Representative plot of calcium transients (CaTs) of cells derived at 10% O_2_. **k**, Representative plot of calcium transients (CaTs) of cells derived at 21% O_2_. **l**, Rise time (s). **m**, Rise speed (fluorescence/s). **n**, Decay time (s). **o**, Decay speed (fluorescence/s). Statistical analysis included Outlier test by ROUT, Normality test by D’Angostino & Pearson test, Unpaired t-test for normal distributed data and Mann-Whitney test for non-normal distributed data. \**P*≤0.05, \*\*\**P*≤0.001, \*\*\*\**P*≤0.0001. **p**, Normalized mRNA levels before derivation and in the early stage of iPS-CM derivation (see Figure S2a) analyzed by qRT-PCR of *MESP1* and *ALDH1A2* (normalized to *PGK1* and *B2M*). Mean, SD, n = 3-4. Statistical analysis included assumption of normality, outlier test by ROUT, and two-way ANOVA followed by Sidak test, \*\*\**P*≤0.001 \*\*\*\**P*≤0.0001.

## Discussion

Based on our observations herein, we propose that sustained low O_2_ levels during iPSC-CM differentiation promote atrial over vCM subtype specification. This outcome was unexpected, as we followed a well-established Wnt-modulation protocol(26) that yielded 70% iPSC-vCM under atmospheric O_2_ (Figure 1f), whereas prior studies(18, 21–23) have shown improved PSC-vCM yield and maturation at lower O_2_ conditions(18, 21–23).

Standard protocols using Wnt modulation typically produce a mix of CM subtypes, including nodal-like cells, ventricular and atrial CM (aCM)(17, 35, 36). During development, CM subtypes arise from distinct progenitors(37): the first heart field (FHF), which forms first(38) and gives rise to vCM and some atrial cells(39–42), and the second heart field (SHF), where posterior SHF (pSHF) gives rise to sinus venosus and aCM(43, 44). Cardiac progenitor specification begins before MESP1 expression(45), with SHF progenitors expressing MESP1 later than FHF(46). We observed differential MESP1 expression by D3 (10- vs. 21% O_2_, Figure S2b), suggesting that O_2_ tension may already influence the cardiac progenitor fate. Similar early progenitor divergence has been reported in other models(47). One study using CHIR99021 reported predominantly FHF-derived vCM in 2D culture but detected SHF markers in another dataset using 3D conditions(9), possibly implicating the physical environment in heart field specification. Another study found that Wnt signaling aligns more closely with SHF than FHF expression profiles(47). In our 10% O_2_ condition, we detected upregulation of atrial markers but expression of both TBX5 (FHF)(8) and NR2F2 (SHF)(47) (Figure 2c), suggesting a mixed population. While these late-stage analyses (D30) may obscure the earlier heart field-specific transcriptional differences, our findings raise the possibility that O_2_ availability influences susceptibility to FHF or SHF specification early during derivation. This was supported by our D3 data, reflecting that 10% O_2_ directs the iPSC-CM into RALDH2+ (*ALDH1A2,* Figure 3p) mesodermal SHF progenitors, whereas atmospheric O_2_ more likely promotes a mesodermal lineage fate with FHF desendants(19).

*In vivo*, mammalian CM experience a range of O_2_ tensions. During cardiac development, CM reside in a relatively hypoxic environment, with approx. 3% O_2_ in developing humans (∼21 mmHg) in the first trimester(10) whereas 5% intramyocardial O_2_ (∼36 mmHg) has been measured in the adult mouse left ventricle(48). After birth there is a marked increase in overall blood oxygenation, from approximately 25-30 mmHg to 75-85 mmHg within minutes(11). In our system, 10% O₂ approximates 76 mmHg and thus exceeds developmental levels and nears adult tissue oxygenation (75-100 mmHg), making the pronounced difference between 10% and 21% O₂ surprising, as 21% O₂ corresponds to 160 mmHg, which is far beyond what is present inside developing cardiac tissue. Yet, whether these approximated O₂ tension in absolute values also reside in the vicinity of the cells in the medium of our system or in general during culture remains speculative, despite that we are in complete control (Figure S1) of the atmosphere in the SCI-TIVE Quad Physoxia glovebox platform exploited. It is possible, that the 2.5–5% O₂ in our system reflects true hypoxia, which could then explain the striking divergence from 10% O₂ with lower iPSC-CM survival as also observed by others(22), despite similarly high Troponin T levels across all groups (Figure 1k). The latter may indeed suggest that CM specification as such is not affected by O_2_ levels, while the CM subtype specification dramatically changes with altered O_2_ levels especially around 10% O_2_. With this, we reject our original hypothesis of enhancing ventricular maturation through simulation of near-physiological O_2_ levels. Some discrepancy exists in the literature regarding the benefits of "low O_2_" during iPSC-vCM derivation. Successful stretches of low O_2_ during vCM derivation from iPSC use 3D cultures in the form of embryoid bodies(19, 22) and/or bioreactor set-ups for derivation and generally finds increased vCM yield and maturation(18, 21). Yet, others have reported that both a low O_2_ pulse and sustained low O_2_ inhibits vCM differentiation(20), and that 3D but not 2D culture benefits from low O_2_(22). Moreover, another research group initially suggested that low O_2_ was beneficial during D0-2 but otherwise dispensable(23) but they have since then cautioned against even short pulses of 5% O_2_(16) based on the promotion of anaerobic glycolysis, an energy pathway CM later abandon in favor of fatty acid oxidation(28). In our protocol, we introduced 10% O₂, prior to metabolic selection and maturation (Figure 1c) and we observed a notable shift in CM identity in a Wnt-modulated iPSC differentiation system. These findings raise the possibility that previously reported inhibitory effects of hypoxia on CM differentiation(16, 20), in part, may reflect the emergence of non-ventricular or otherwise unintended cardiac subtypes.

Specification of aCM from pSHF and iPSC is generally thought to be dependent on retinoic acid (RA) signaling during cardiac mesoderm formation(49–51), and reduced Activin A and BMP4 also promote an atrial fate(19). Atrial fibrillation (AF) is the most common sustained cardiac arrythmia, however animal models are inherently limited in their ability to replicate AF largely due to the absence of naturally occurring AF(52). As a result, iPSC-derived aCM have become valuable tools for AF disease modeling(53), personalized medicine and drug screening(54). However, still challenges with cellular immaturity and capturing the atrial/AF phenotype exist(55, 56). While our study was designed to enhance iPSC-vCM specification, we acknowledge that the resulting observed atrial-like populations at present may not represent fully matured aCM and there are limitations accordingly. Nonetheless, cells differentiated at 10% O₂ without RA treatment herein, exhibit features consistent with an aCM identity, to a degree comparable to the ventricular characteristics seen in cells derived at 21% O₂ (Figure 2f). Future studies should refine the aCM 10% O₂ differentiation protocol applied here, potentially integrating strategies already established for atrial lineage induction, such as RA(19). An interesting subject, but also a point of concern for future iPSC-vCM studies, including manufacturing, is the possibility that O_2_ gradients within 3D spheroid iPSC-vCM cultures may facilitate unwanted differential cardiac subtype specification, with a hypoxic core favoring atrial specification, whereas the normoxic surface promotes a ventricular lineage. Likewise, it is possible that also other organ differentiation systems involving organoid formation may be similarly affected by the O_2_ tension which could harness their diagnostic or regenerative uses.

A notable limitation of the present study is the reliance on a single, albeit widely used, iPSC line which may constrain transferability due to potential line-specific genetic or epigenetic effects, and we want to emphasize that our findings should be interpreted within the context of this specific genetic background. Nevertheless, across four independent experiments we consistently observed an atrial phenotype and believe this unexpected observation could be valuable for planning experiment where similar oxygen-regulated effects might arise. Further studies using multiple iPSC lines with diverse genetic backgrounds will be essential to determine whether this effect extends beyond the WTC-11 line. In conclusion, our study using sustained and tightly controlled O_2_ levels during iPSC-CM derivation unraveled that more physiological levels of "low" O_2_ at 10% O_2_ limits vCM subtype specification but may serve as an attractive approach to refine atrial CM specification from iPSCs. Further investigation into the microenvironmental O_2_ cues thus may enhance the relevant fields abilities to direct more precise iPSC-CM subtype-differentiation to benefit their use for diagnostic-, cytotoxic-, and regenerative purposes.

## Methods

### Induced pluripotent stem cell culture and cardiomyocyte derivation

The WTC-MYL2v-GFP iPSC line (passage 37), with a mono-allelic mEGFP tag to the c-term of MYL2(57) (AICS-0060-027) was developed at the Allen Institute for Cell Science (allencell.org/cell-catalog) and available through Coriell, who performed the authentication and mycoplasma and sterility screening. Maintenance and differentiation were overall performed as previously described(26) and passages up until p59 were used for experiments. Undifferentiated iPSCs were maintained on approximately 10 µg/cm^2^ growth factor reduced Matrigel (Corning, 356231)-coated surfaces at 21% O_2_, 37 °C and 5% CO_2_ in mTeSR+ medium (Stem Cell Technologies, 100-0274), passaged with Accutase (Thermo Fisher, A11105-01) and seeded at a density of 50,000 cells per well in TPP 6-well plates (5,539 cells/cm^2^). All passages were performed with 10 µM Rock inhibitor Y27632 2HCl (Ri) (Selleckchem, S1049) added to the medium and a subsequent replenish of media without Ri after 24 h. For revival of iPSC, vials were quickly thawed, and cells were dropwise transferred into 9 mL pre-warmed media, and the vial was rinsed in 1 mL before centrifugation, resuspension in warmed mTeSR+/Ri and seeding in 6-well plates. Cells were frozen by diluting cell suspension in mTeSR+/Ri 1:1 in freezing medium consisting of mTeSR+/Ri and 20% DMSO. For cultivation at 2.5-, 5-, and 10% O_2_ tensions, iPSCs were transferred to pre-equilibrated SCI-TIVE Quad Physoxia closed glovebox workbenches (Baker-Ruskinn, serial number: 0118NSC01), which ensured continuous monitoring and control over culture conditions (Humidity, gas composition (O_2_/CO_2_/N_2_) and temperature (Figure S1) during iPSC-CM derivation. All media were supplemented with 1% Penicillin/streptomycin (PS) (5,000 U/ml, 15070-063, Gibco). For iPSC-CM derivation (Figure 1a), cells were counted (ChemoMetec, NC-200/NC-202) and 20,000 iPSC/cm^2^ were plated in mTeSR+Ri three days prior (derivation day -3 (D-3)) to derivation start D0. For CM derivation at 2.5-, 5-, and 10% O_2_ tensions, iPSCs were transferred to SCI-TIVE Quad Physoxia closed glovebox workbenches on D-2 and all further actions hereafter up until fixation were performed inside the glovebox to avoid exposure to ambient air. CM differentiation was performed using modulators of the wnt pathway contained in B27 medium with or without insulin as previously described(26). Metabolic selection(28) was undertaken for four days before maturation and fixation on day 30.

### RNA, cDNA and qRT-PCR

Quantitative reverse transcriptase PCR (qRT-PCR) was performed as previously described(58). Cells were lysed in TriReagent (Invitrogen, 15596018) whereafter RNA was isolated with Polyacryl carrier (PC 152; MRC), BCP (1-Bromo-3-Chloro-Propane) (Sigma-Aldrich, B9673) and 2-propanol (Sigma-Aldrich, I9516), and lastly rinsed in 75% ice cold ethanol. After resolving in nuclease-free water, the concentration was measured via nano-drop(29).

cDNA was generated using the High-Capacity cDNA Reverse Transcriptase kit (Applied Biosystems, 4368813) according to manufacturer’s protocol. For qRT-PCR, each sample was mixed to a total volume of 10 µl, containing a mixture of 2 ng cDNA (template), 0.3 µM forward and reverse primers (Supplementary Table 1), and Power SYBRGreen PCR Master Mix (Applied Biosystems, 4367659). Each sample was analyzed in technical triplicates. Samples were run at a 7900HT Fast Real-time PCR system (Applied Biosystems): 10 min holding at 95°C, 40 cycles of 15 sec denaturation at 94°C, 30 sec of annealing at 60°C and 30 sec of elongation at 72°C. Melting curves and amplification efficacy were used for quality assessment. For *ALDH1A2* a TaqMan assay (Thermo Fisher, Hs00180254_m1) was used according to the manufacture recommendations. For all assays normalization was performed against multiple stably expressed endogenous genes as determined by the geNorm and qBase Plus 3.2 platform (Biogazelle).

## Flow cytometry

Cells were dissociated and fixed in 2.5% NBF diluted in Hanks Balanced Salt Solution (HBSS) (EuroClone, ECM0512L) containing 5% FBS (Sigma-Aldrich, F0804-500ML) and 1% PS (DE17-602E; Lonza) (HBSS/5%FBS/1%PS) for 15 min. After three washes in HBSS/5%FBS/1%PS, the cells were stored at 4°C in HBSS/5%FBS/1%PS containing 0.05%NaN_3_. For staining, fixed cells were permeabilized in PBS with 1% BSA and 0.1% TX100 (PBS/1%BSA/0.1%TX100) and stained with mouse anti-Troponin T (Life technologies, MA116687, 1:500) and rabbit anti-GFP (Abcam, ab290, 1:500) for 1h on ice while shaking. After washing twice in PBS/1% BSA/0.1% TX100, the cells were incubated with secondary antibodies 488-donkey anti-rabbit (Invitrogen, A21206, 1:200) and 555-donkey anti-mouse (Invitrogen, A31570, 1:200) for 30 min on ice while shaking. After 2 final washings in PBS/1%BSA/0.1%TX100, Hoechst (Sigma, 33342, 1:500) were added to the cells, and they were analyzed on a LSRII flow cytometer (BD Biosciences). Analysis was performed using the FACSDiva software version 8.0.1, by serially gating according to size and granularity in forward-versus side scatter and then for incorporation of Hoechst before sub-fractionated into an iPSC-CM pool based on the presence of Troponin T and then vCM based on GFP expression. Ploidy was analyzed by Hoechst subpopulations, 2n, 4n and >4n, of troponin T^+^ iPSC-CM. The percentage and geometric mean were determined.

## Immunocytochemistry

Immunocytochemistry was performed according to a previous report(58). After a brief wash in phosphate buffered saline (PBS), cultured cells were fixed in 10% Neutral Buffered Formalin (NBF) (HT501128, Sigma-Aldrich) for 10 min, washed three times in PBS, and stored in PBS containing 0.05% NaN_3_ at 4°C in the dark. Samples were permeabilized in Tris-Buffered Saline (TBS) with 0.3%Triton-x-100 (Sigma-Aldrich, TX100) for 10 min and then blocked in TBS containing 2% BSA for 20 min before overnight incubation with primary antibodies diluted in TBS with 1% BSA at 4°C while gently shaking. The antibodies used were: Rabbit anti-GFP (1:500), mouse anti-Cardiac Troponin T antibody clone 1C11 (1:500), rabbit anti-GATA4 (Santa Cruz Biotechnologies, sc-9053, 1:50), rabbit anti-NKX2.5 (Santa Cruz Biotechnologies, sc-14033, 1:100), mouse anti-Tropomyosin clone CH1 (1:500, T9283, Sigma), and mouse anti-sarcomeric actinin (Sigma, A7811, 1:200). After three washes in TBS, cells were incubated with appropriate secondary antibodies: 488-donkey anti-rabbit (Invitrogen, A21206, 1:200), 555-donkey anti-rabbit (Invitrogen, A31572, 1:200), or 647-donkey anti-mouse (Invitrogen, A31571, 1:200) for 1h at room temperature while shaking in the dark. DAPI (Vector Lab, Vectashield, H-1200) was added to the stained cells after three final TBS washes and investigated by microscopy on a Leica DMI 4000 B microscope with a Leica CTR4000 illuminator and a Leica DFC300FX/DFC 340 FX camera. Camera settings and image editing were kept identical between samples and controls. Quantification of immunostained images was performed using the Fiji 2 (ImageJ2) version 2.14.0/1.54j.

## ScRNA-seq

ScRNA-seq was performed as previously described(29). To avoid RNA degradation, the environment and all reagents were kept strictly RNAse free. This protocol was tested beforehand for cell clumping by microscopy and for RNA preservation through determination of the RNA integrity number (RIN) on Agilent 2100 Bioanalyzer (Agilent Technologies) with Agilent RNA 6000 Nano Kit (Agilent Technologies)(59). For scRNA-seq iPSC-CM generated under the four different O_2_ paradigms were rinsed in warmed RNAse free PBS and dissociated with pre-warmed trypsin/EDTA (0.05%). Detached cells were diluted in 2 mL RNAse free PBS with 1% RNAse free BSA (VWR, 0332-100G) (PBS/1%BSA) and gently pipetted up and down to avoid clumps. The wells were rinsed in 1 ml RNAse free PBS to collect all cells. After centrifugation at 300 g for 5 min at 4°C, the pelleted cells were washed in 1 ml chilled RNAse free PBS before methanol fixation for 15 min on ice. Rehydration was performed to reconstitute the RNA to its original state(29).

Libraries were prepared using the Chromium Next GEM Single Cell 3’ GEM Kit v3 16 runs (10X Genomics, PN-1000123), following the manufactures instructions. The cDNA libraries were sequenced using the Illumina NovaSeq 6000 System (Illumina inc., 20012850). After sequencing, data was processed using the CellRanger suite (v. 7.0.1). Reads were demultiplexed using CellRanger mkref and subsequently aligned to the GrCH38 reference genome and quantified using CellRanger count. Samples were sequenced at a mean depth of 319 million reads, resulting in an average depth of 33,771 reads per cell. Quantified reads were further processed in R (v. 4.4.1) using the package Seurat (v. 5.1.0). The quality metrics of the scRNA-seq dataset have previously been reported partly when validating the fixation protocol(29). A detailed quality assessment of all samples is provided in supplemental Fig. S3. Quality checked data was filtered by excluding low-quality cells wherein less than 300 genes were detected and more than 15% of the reads mapped to mtRNA. Low detectable genes that were found in less than 3 cells were likewise excluded. A total of 28,689 quality passed cells across the 4 samples were further subjected to normalization, and dimensional reduction. In short, samples were handled as a single object, normalized using the Seurat log-normalization procedure, scaled, and the top 3,000 variable genes were extracted, and applied for dimensional reduction using UMAP. The low dimensional space was clustered using Louvain clustering at a resolution of 0.3. For comparison of iPSC-CM with adult cardiomyocytes from atrial or ventricular chambers, we acquired data from Litviňuková et al.(32). FASTQ-files from libraries of the myocardium including, left or right ventricle and left or right atrium were downloaded from the European Nucleotide Archive (ENA, https://www.ebi.ac.uk/ena/browser/view/PRJEB39602). Data was aligned and quantified as described above. For integration, cells that had more than 1500 genes detected were included in the analysis. Filtered cells were normalized using SCTransform with the variance stabilization method v1 applied(60). The top 500 most variable genes were used for dimensional reduction. To correct for experimental variability between adult CM and iPSC-CM Harmony integration(61) were used. Harmony corrected data was then dimensionally reduced as performed and described above.

## Calcium imaging

For cytosolic calcium transient recordings, D24 iPSC-CM derived at 10- or 21% O_2_ were replated at low density in 6-well plates and cultured at 10- or 21% O_2_ until D30 in FAM. To visualize the calcium transients, iPSC-CM were incubated with 5 µM Rhod-2 AM fluorescent calcium indicator (Thermo Fisher, R1244) in Tyrode’s solution (0.02% Pluronic F-127, 140 mM NaCl, 5.4 mM KCl, 1.8 mM CaCl, 1 mM MgCl, 10 mM HEPES, 10 mM glucose and pH adjusted to 7.4) for 20 min at 37 °C and 5% CO_2_. Cells were finally washed and kept in Tyrode’s solution for imaging at room temperature. Fluorescent images were recorded using an Olympus FV1000 MPE confocal microscope with a UMPLFL 10x W NA: 0.30 objective. Rhod-2 was excited at 559 nm, and fluorescence was collected at 577 nm. The scan area was 128x128 pixels, and images were recorded every 0.188 sec for 60 sec. The temporal profile of Rhod2 fluorescence was extracted from the cytosolic region of 5 randomly selected cells using Fiji ImageJ and analyzed using SciPy (find_peaks, peak_widths) in Python 3.

## Statistics

All data are presented in the figures, and each analysis consisted of at least three independent experiments designated n, and for some experiments like qRT-PCR each n consisted of three technical replicates. For cell line experiments, independent experiments (n) were separated by at least two passages. If relevant, exclusion of outliers was performed by ROUT, whereas normal distribution was tested by D’Agostino & Pearson test. Statistical significance of the difference between means was determined by appropriate tests as indicated in the figure legends or text. The GraphPad Prism version 10.5.0 was used for all statistical analysis. We considered significance levels α = 0.05 for identifying significant results marked by asterisks *P<0.05, **P<0.01, ***P<0.001, ****P<0.0001, ns (not significant).

## Declarations

### Ethics approval and consent to participate

N/A

## Consent for publication

N/A

## Data availability

ScRNA-seq data that supports the findings of this study have been deposited in the Gene Expression Omnibus (GEO) under the accession code GSE299973. All other data supporting the findings of this study are available from the corresponding author on a reasonable request.

## Author contributions

SBM, FAB, AKST, DGE, JHL: Collection of data, data analysis and interpretation, manuscript writing, and final approval of manuscript. PBH: Collection of data and final approval of manuscript. CHJ, ENYP, PS: Conception and design, data interpretation, and final approval of manuscript. DCA: Conception and design, collection of data, data analysis and interpretation, manuscript writing, final approval of manuscript, and financial support.

## Acknowledgements

We would like to thank Adrian Calmar and Claus Jespersen (Andersen group, Odense University Hospital/University of Southern Denmark) for excellent technical assistance on this study, Christina D. Fenger (Amplexa Genetics, Odense) for cDNA library preparation and assistance of sequencing, and Mark Burton (Department of Clinical Genetics, Odense University Hospital) for assisting with guidance and analysis of raw sequencing data.

