## Supplemental material for "Sustained 10% Oxygen Promotes Atrial Rather Than Ventricular Specification During Human iPSC-Cardiomyocyte Differentiation"

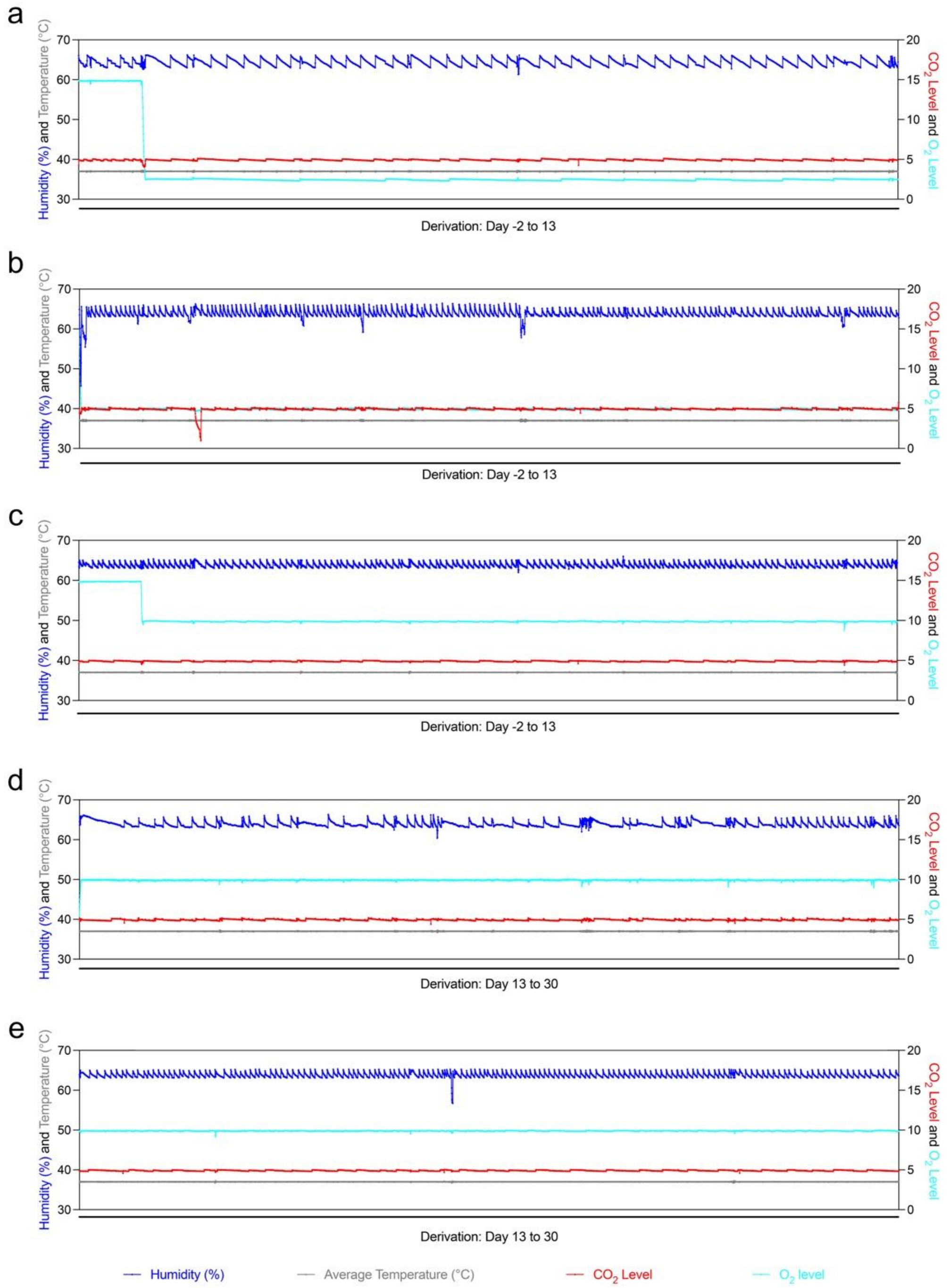

**Figure S1. Humidity, temperature, and gas composition in SCI-TIVE Quad Physoxia closed glovebox workbenches during iPSC-CM derivation. a-c**, Humidity, temperature, $CO_2$ and $O_2$-tension during derivation at 2.5-, 5-, and 10% $O_2$, respectively, at derivation day (D) D-2 to D13. **d-e**, Humidity, temperature, $CO_2$ and $O_2$-tension in the two chambers used during derivation at 10 % $O_2$ at D13 to D30.

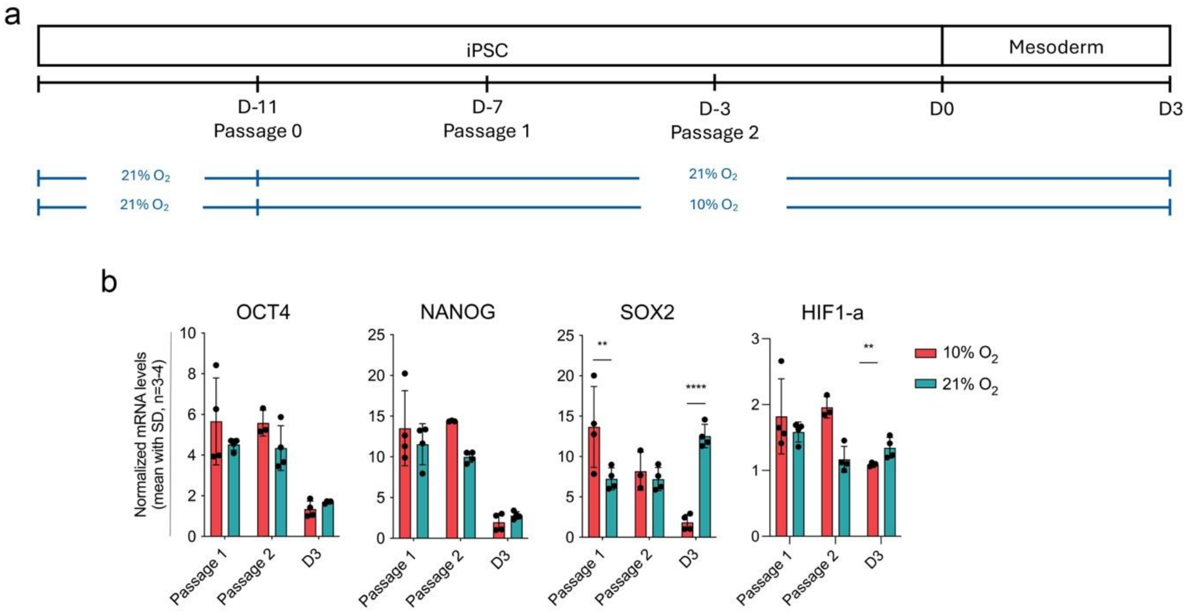

**Figure S2. Effects of 10- and 21% $O_2$ tensions on pluripotency of iPSCs before derivation and in early stages of iPSC-CM derivation. a**, Schematic overview of $O_2$ tensions before derivation and in the early stage of iPSC-CM derivation. **b**, Normalized mRNA levels analyzed by qRT-PCR of *OCT4, NANOG, SOX2* (pluripotency markers) and *HIF-1α* (hypoxia marker) for the indicated passage- and derivation days (normalized to *PKG1* and *B2M*). Mean, SD, n = 3-4. Statistical analysis included assumption of normality, outlier test by ROUT, and two-way ANOVA, $^{**}P \leq 0.01$ $^{****}P \leq 0.0001$.

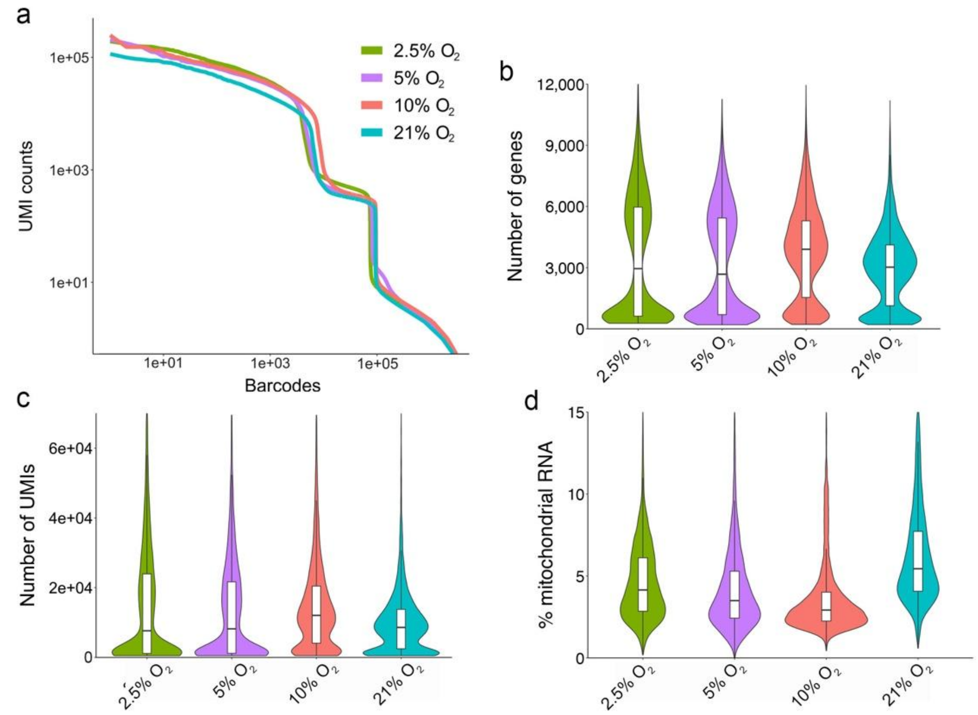

e

| # of cells | 2.5% $O_2$ | 5% $O_2$ | 10% $O_2$ | 21% $O_2$ |
|---|---|---|---|---|
| Total | 5431 | 6449 | 9611 | 7198 |
| ACTC1 | 5356 | 5443 | 9571 | 7155 |
| TNNT2 | 5375 | 5403 | 9594 | 7197 |
| TNNC1 | 5378 | 5400 | 9595 | 7193 |
| Filtered | 5241 | 5190 | 9538 | 7180 |

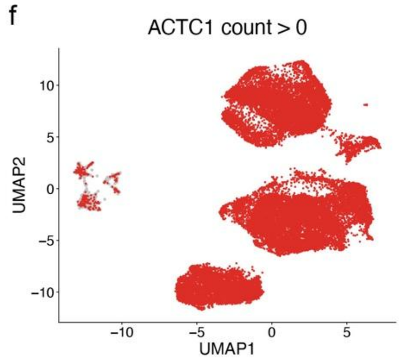

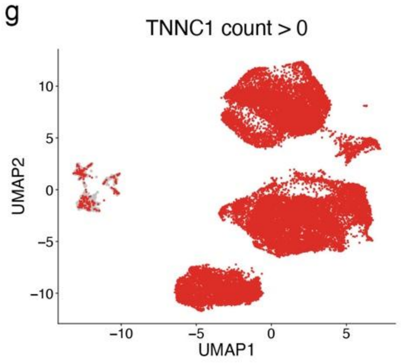

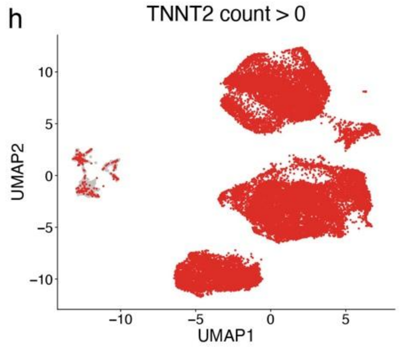

**Figure S3. scRNA-seq quality assessment and iPSC-CM purity. a-d**, Quality analyses of scRNA-seq iPSC-CMs data (2.5-, 5-, 10-, and 21% $O_2$). **a**, Barcode rank plot (knee-plot) showing log scaled barcodes ranked by the number of Unique Molecular Identifier (UMI) vs. log scaled UMI counts. **b**, Violin plot showing "Median Genes per Cell". **c**, "Median UMI Counts per Cell". **d**, Percentages of mitochondrial transcripts. **e**, Table showing the total number of cells prior and after analytic CM filtering according to the expression of CM marker genes *ACTC1*, *TNNT2*, and *TNNC1*. **f-h**, Uniform Mainfold Approximation Projection (UMAP) plots showing the expression of the CM markers **f**, *ACTC1,* **g**, *TNNC1*, and **h**, *TNNT2* prior to filtering (red: positive cells, grey: negative cells).

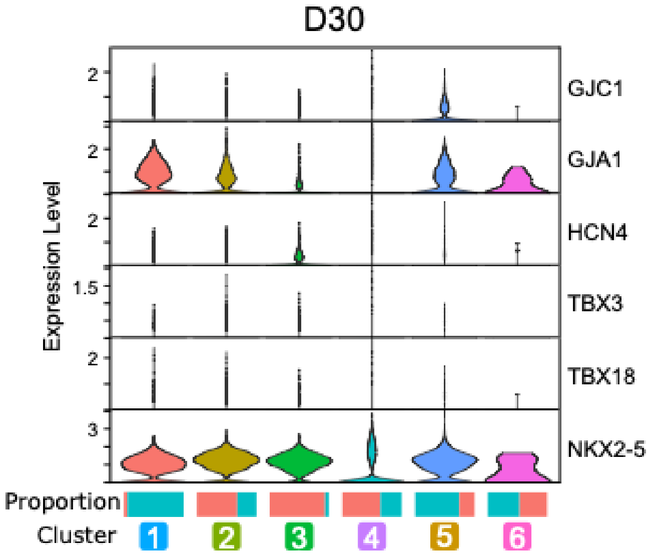

**Figure S4. Neither 10- nor 21% $O_2$ drives iPSC pacemaker cell specification.** Violin plots visualizing normalized expression of known positive (*GJC1, HCN4, TBX3, TBX18*) and negative (*GJA1* (also known as Cx43), *NKX.2-5*) marker genes for pacemaker cells. Proportions of 10- and 21% $O_2$-derived iPSC-CMs for each cluster are visualized as bar plots below the plot (Refer to Figure 2 a-c in main figures).

**Supplementary Table 1. qPCR primers - 5' to 3'**

| | Forward | Reverse |
|---|---|---|
| OCT4 | CAGTGCCCGAAACCCACAC | GGAGACCCAGCAGCCTCAAA |
| TNNT2 | GACAGAGCGGAAAAGTGGGA | GCGGGTCTTGGAGACTTTCT |
| MYH6 | GATAGAGAGACTCCTGCGGC | GGTTCTCCCGATCTGTCAGC |
| MYH7 | TCCCCACCATCTCTTTCCCTCGTA | TCCTGACACTGCCCCTGAACCA |
| MYL2 | TGTCCCTACCTTGTCTGTTAGCCA | ATTGGAACATGGCCTCTGGATGGA |
| SCN5A | TCTTCACAGGCGAGTGTATTG | GACAACCACGAAGTCGAAGATA |
| SIRPA | ACCTGGCTCAGGCTAGTTCCAAAT | TGTGCACACGTATGTGCTGTCTCT |
| FABP3 | CACTCGCACTTATGAGAAAGA | AGGAAGAAATGAGGCAATGT |
| CKMT2 | GGAGAGAGGCCAAGATATTAAG | CGTACCAGCAGACAGATTATT |
| NANOG | CGAAGAATAGCAATGGTGTGACG | TTCCAAAGCAGCCTCCAAGTC |
| SOX2 | CCCTGTGGTTACCTTTTCCT | AGTGCTGGGACATGTGAAGT |
| MESP1 | TCGAAGTGGTTCCTTGGCAGAC | CCTCCTGCTTGCCTCAAAGTGTC |
| HIF-a | CCAGCAGACTCAAATACAAGAACC | TGTATGTGGGTAGGAGATGGAGAT |
| hGAPDH | GCCACATCGCTCAGACACCATGG | TCCCGTTCTCAGCCTTGACGGT |
| hPgk1 | GTCGGCTCCCTCGTTGACCGAA | GGGACAGCAGCCTTAATCCTCTGGT |
| hB2M | GCCTGCCGTGTGAACCATGTGA | ATGCGGCATCTTCAAACCTCCATGA |
| hHprt-1 | GGCTCCGTTATGGCGACCCG | CCCCTTGAGCACACAGAGGGCTA |
